# Hydrogenotrophic methanogens of the mammalian gut: functionally similar, thermodynamically different - A modelling approach

**DOI:** 10.1101/445171

**Authors:** Rafael Muñoz-Tamayo, Milka Popova, Maxence Tillier, Diego P. Morgavi, Jean-Pierre Morel, Gérard Fonty, Nicole Morel-Desrosiers

## Abstract

Methanogenic archaea occupy a functionally important niche in the gut microbial ecosystem of mammals. Our purpose was to quantitatively characterize the dynamics of methanogenesis by integrating microbiology, thermodynamics and mathematical modelling. For that, *in vitro* growth experiments were performed with pure cultures of key methanogens from the human and ruminant gut, namely *Methanobrevibacter smithii, Methanobrevibacter ruminantium* and *Methanobacterium formicium*. Microcalorimetric experiments were performed to quantify the methanogenesis heat flux. We constructed an energetic-based mathematical model of methanogenesis. Our model captured efficiently the dynamics of methanogenesis with concordance correlation coefficients of 0.94 for CO_2_, 0.99 for H_2_ and 0.97 for CH_4_. Together, experimental data and model enabled us to quantify metabolism kinetics and energetic patterns that were specific and distinct for each species despite their use of analogous methane-producing pathways. Then, we tested *in silico* the interactions between these methanogens under an *in vivo* simulation scenario using a theoretical modelling exercise. *In silico* simulations suggest that the classical competitive exclusion principle is inapplicable to gut ecosystems and that kinetic information alone cannot explain gut ecological aspects such as microbial coexistence. We suggest that ecological models of gut ecosystems require the integration of microbial kinetics with nonlinear behaviours related to spatial and temporal variations taking place in mammalian guts. Our work provides novel information on the thermodynamics and dynamics of methanogens. This understanding will be useful to construct new gut models with enhanced prediction capabilities and could have practical applications for promoting gut health in mammals and mitigating ruminant methane emissions.

## Introduction

Methanogenic archaea inhabit the gastro-intestinal tract of mammals where they have established syntrophic interactions within the microbial community (1–3) playing a critical role in the energy balance of the host (4,5). In the human gut microbiota, the implication of methanogens in host homeostasis or diseases is poorly studied, but of growing interest (6). *Methanobrevibacter smithii* (accounting for 94% of the methanogen population) and *Methanosphaera stadtmanae* are specifically recognized by the human innate immune system and contribute to the activation of the adaptive immune response (7). Decreased abundance of *M. smithii* was reported in inflammatory bowel disease patients (8), and it has been suggested that methanogens may contribute to obesity (9). In the rumen, the methanogens community is more diverse though still dominated by *Methanobrevibacter* spp., followed by *Methanomicrobium* spp., *Methanobacterium* spp. (10) and *Methanomassillicoccus* spp (11). However, the proportion of these taxa could vary largely, with *Methanomicrobium mobile* and *Methanobacterium formicium* being reported as major methanogens in grazing cattle (12). Though methanogens in the rumen are essential for the optimal functioning of the ecosystem (by providing final electron acceptors), the methane they produce is emitted by the host animal, contributing to global greenhouse gas (GHG) emissions. The livestock sector is responsible for 14.5% of the anthropogenic GHG emissions, of which enteric methane from ruminants contributes to more than 40% of total emissions (13). Methanogens can be separated in two groups based on the presence or absence of cytochromes, which are membrane-associated electron transfer proteins providing energetic advantages for microbial growth (14). However, in the gastrointestinal tract of mammals, the only methanogens with cytochromes are the *Methanosarcinales* which are minor members of the community (4,15). Major rumen methanogens (16) and the dominant human archaeon *M. smithii* (17), are hydrogenotrophic archaea without cytochrome. Cytochrome-lacking methanogens exhibit lower growth yields than archaea with cytochromes (14). However, this apparent energetic disadvantage has been counterbalanced by a greater adaptation to the environmental conditions prevailing in the gastrointestinal tract (15), and by the establishment of syntrophic interactions with feed fermenting microbes. This syntrophic cooperation centred on hydrogen allows the anaerobic reactions of substrate conversion to proceed close to the thermodynamic equilibrium (18,19) (that is with Gibbs free energy change close to zero).

To our knowledge, the impact of thermodynamics on human gut metabolism has been poorly addressed in existing mathematical models (20–23). For the rumen, thermodynamic principles have been incorporated already into mathematical research frameworks because of their important role in feed fermentation, (24–29). Despite these relevant efforts, much work remains to be conducted for attaining a predictive thermodynamic-based model that allows for quantitative assessment of the impact of the thermodynamics on fermentation dynamics. Theoretical frameworks have been developed to calculate stoichiometric and energetic balances of microbial growth from the specification of the anabolic and catabolic reactions of microbial metabolism (30,31), and advances have been done to link thermodynamics to kinetics (32–34). These works constitute a solid basis for tackling the thermodynamic modelling of gut metabolism. In this respect, new knowledge on the extent of methanogenesis could help to improve existing gut models. Accordingly, our purpose was to quantitatively characterize the dynamics of hydrogen utilization, methane production, growth and heat flux of three hydrogenotrophic gut methanogens by integrating microbiology, thermodynamics, and mathematical modelling. We investigated the rate and extent of methanogenesis by performing *in vitro* experiments with three methanogenic species representing major human and ruminant genera: *M. smithii, M. formicium* and *Methanobrevibacter ruminantium*. To interpret and get the most out of the resulting data, a mathematical model with thermodynamic basis was developed to describe the dynamics of the methanogenesis. Our findings allowed to quantify metabolism kinetics and energetic patterns that were specific and distinct for each species despite their use of analogous methane-producing pathways and their common belonging to the group of cytochrome-lacking methanogenic archaea.

## Material and Methods

### *In vitro* growth experiments

#### Archaeal strains and growth media

Archaeal strains used in the study were *M. ruminantium* M1 (DSM 1093), *M. smithii* PS (type strain DSM 861), and *M. formicium* MF (type strain DSM 1535). The growth media was prepared as previously described (35) and composition is summarized in Table S1 of the Supplementary material.

#### Experimental design and measures

Starter cultures were grown until reaching optical density at 660 nm (OD_660_) of 0.400 ± 0.030. Optical density was measured on a Jenway spectrophotometer (Bibby Scientific). Then, 0.6 ml were used to inoculate one experimental tube. Commercially prepared high purity H_2_/CO_2_ (80%/20%) gas mix was added to inoculated tubes by flushing for 1 min at 2.5 Pa. Mean initial OD_660_ and pressure values are summarized in Table S2 of the Supplementary material. Growth kinetics for each strain were followed over 72 h. The experiment was repeated twice. Each kinetics study started with 40 tubes inoculated at the same time. At a given time point (3 h, 5 h, 6.5 h, 24 h, 26 h, 30 h, 47h, 72 h post inoculation), two tubes with similar OD_660_ values were sampled. The tubes were used for measuring gas parameters: pressure was measured using a manometer and composition of the gas phase was analysed by gas chromatography on a Micro GC 3000A (Agilent Technologies, France). One of the tubes was centrifuged 10 min at 13 000 g. The microbial pellet was weighed and stored at −20°C in 2 ml screw-cap tubes containing 0.4 g of sterile zirconia beads (0.3 g of 1 mm and 0.1 g of 0.5 mm).

#### DNA extraction and qPCR quantification of 16S rRNA genes

One ml of lysis buffer (50mM NaCl, 50 mM TrisHCl pH 7.6, 50 mM EDTA, 5 *%* SDS) was added directly to the frozen microbial pellet before homogenizing for 2 × 30 s at 5100 tours/min in a Precellys bead-beater (Bertin Instruments). Samples were centrifuged for 3 min at 14 000 g and the liquid phase transferred to a new tube before adding 600 μl of phenol–chloroform–3-methyl-1-butanol (25:24:1) solution. After centrifugation at 14 000 g for 3 min, the aqueous phase was transferred to a fresh tube and 500 μl of chloroform were added. The chloroform-washing step was repeated twice with centrifugation at 14000 g for 3 min between steps. The final volume of the aqueous phase was measured and DNA precipitation was initiated by adding 70% of the volume of isopropanol 100% and 10% of the volume of sodium acetate 3M. Sedimentation at 14 000 g for 30 min was again performed and the resulting DNA pellet was washed with 500 μl of ethanol 70% and dissolved in 50μl of molecular biology quality water. The extraction yield was checked on a Nanodrop 1000 Spectrophotometer (Thermo Fisher Scientific, France) and extracts run on a FlashGel System (Lonza, Rockland, Inc) to check integrity.

Copies of 16S rRNA genes were quantified using a qPCR approach. Primers used are those of Ohene-Adjei et al (36); reaction assay and temperature cycles were as described previously (37). Triplicate qPCR quantification was performed on 20 ng of extracted DNA. Amplifications were carried out using SYBR Premix Ex Taq (TaKaRa Bio Inc., Otsu, Japan) on a StepOne system (Applied Biosystems, Courtabeuf, France). Absolute quantification involved the use of standard curves that had been prepared with gDNA of *Methanobrevibacter ruminantium* DSM 1093. PCR efficiency was of 103%. Results were expressed as copy numbers per ng of extracted DNA per g of microbial pellet. *M. smithii* and *M. ruminantium* strains used in this study possess two copies of 16S rRNA genes in their genomes. The number of cells was computed by dividing 16S copy numbers by 2.

#### Microcalo rimetry

Microcalorimetric experiments were performed to determine the heat flux pattern of each methanogen. Metabolic activity and microbial growth were monitored by using isothermal calorimeters of the heat-conduction type (A TAM III, TA Instruments, France) equipped with two multicalorimeters, each holding six independent minicalorimeters, allowed continuous and simultaneous recording as a function of time of the heat flux produced by 12 samples. The bath temperature was set at 39°C; its long-term stability was better than ± 1×10^-4^°C over 24h. Each minicalorimeter was electrically calibrated. The specific disposable 4 mL microcalorimetric glass ampoules capped with butyl rubber stoppers and sealed with aluminium crimps were filled with 1.75 mL of Balch growth media and overpressed with 2.5 Pa of H_2_/CO_2_ 80%/20% gas mixture for 30 s. There was no significant difference in pressure at the beginning of the study. They were sterilized by autoclave and stored at 39°C until the beginning of the microcalorimetric measurements. Actively growing cultures of methanogens (OD_660_ of 0.280±0.030 for *M. smithii*, 0.271±0.078 for *M. ruminantium* and 0.142±0.042 for *M. formicium*) were stored at −20°C in order to diminish microbial activity before inoculation. Cultures were thawed for 30 min at ambient temperature and inoculation was carried out by injecting 0.25 mL of the culture through the septum of the overpressed microcalorimetric ampoules just before inserting them into the minicalorimeters. Samples took about two hours to reach the bath temperature and yield a stable zero baseline. Blank experiments were also carried out by inserting ampoules that were not inoculated and, as expected, no heat flux was observed confirming the medium sterility. Each experiment was repeated thrice.

The heat flux 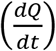 also called thermal power output *P*, was measured for each methanogen and blank samples with a precision ≥ 0.2 μW. The heat flux data of each sample were collected every 5 minutes during more than 10 days. The total heat Q was obtained by integrating the overall heat flux time curve using the TAM Assistant Software and its integrating function (TA Instruments, France).

Classically, the heat flux-time curve for a growing culture starts like the S-shaped biomass curve (a lag phase followed by an exponential growth phase) but differs beyond the growth phase, the heat flux being then modulated by transition periods (38). Heat flux data can be used to infer the microbial growth rate constant provided the existence of a correlation between isothermal microcalorimetry data and microbiological data (e.g., cell counts) at early growth (39). During the exponential growth phase, microbial growth follows a first-order kinetics defined by the specific growth rate constant μ_c_ (h^-1^). Analogously, the heat flux follows an exponential behaviour determined by the parameter μ_c_ as described by (38,39).

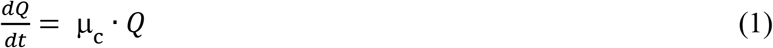

The growth rate constant μ_c_ can be determined by fitting the exponential part of the heat flux-time curve using the fitting function of the TAM Assistant Software. In our case study, careful selection of the exponential phase of heat flux dynamics was performed to provide a reliable estimation of the maximum growth rate constant from calorimetric data.

### Mathematical model development

#### Modelling *in vitro* methanogenesis

The process of *in vitro* methanogenesis is depicted in Figure 1. The H_2_/CO_2_ mixture in the gas phase diffuses to the liquid phase. The H_2_ and CO_2_ in the liquid phase are further utilized by the pure culture to produce CH_4_. Methane in the liquid phase diffuses to the gas phase.

**Figure 1.**
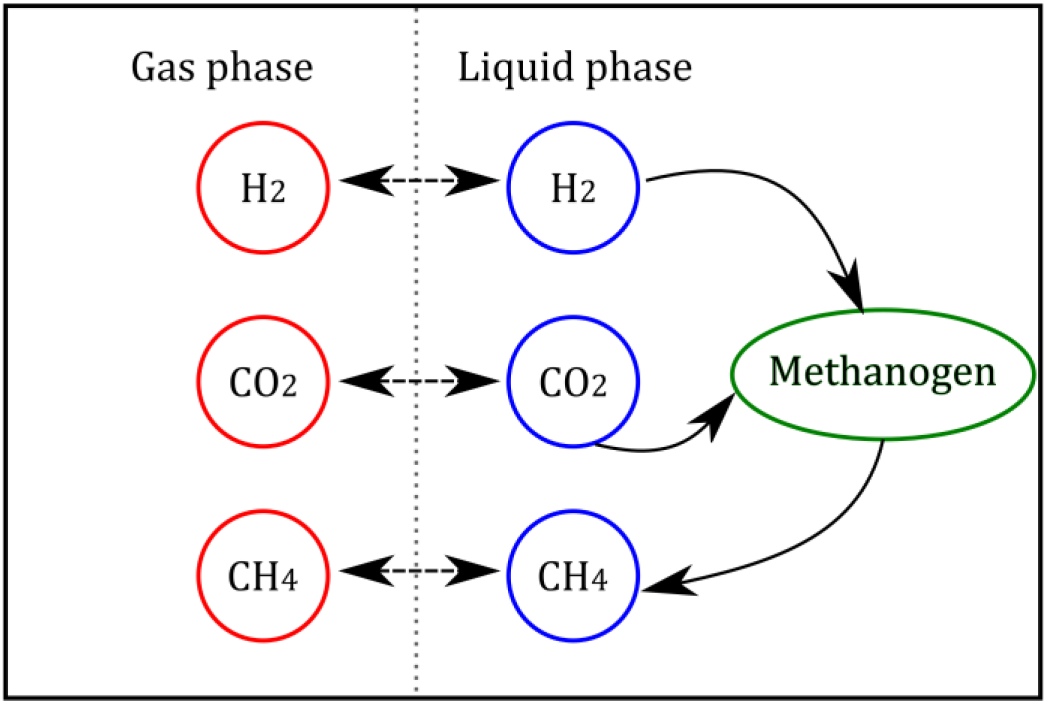
Schematics of the *in vitro* methanogenesis process by hydrogenotrophic methanogens. Double arrows represent fluxes due to liquid-gas transfer, simple arrows represent metabolic fluxes.

Model construction was inspired by our previous dynamic models of human gut (20) and rumen *in vitro* fermentation (40) followed by certain simplifications. The model considers the liquid-gas transfer of carbon dioxide. Due to the low solubility of hydrogen and methane (41), the concentration of these two gases in the liquid phase was not modelled. We assumed that the dynamics of concentrations in the gas phase are determined by kinetic rate of the methanogenesis. To incorporate thermodynamic information, instead of using the Monod equation in the original formulation, we used the kinetic rate function proposed by Desmond-Le Quéméner and Bouchez (33). The resulting model is described by the following ordinary differential equations

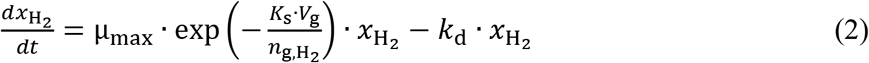

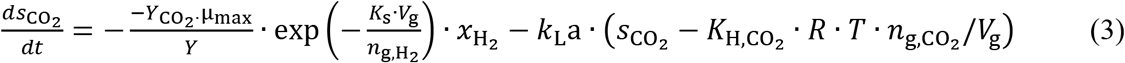

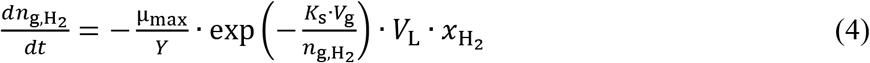

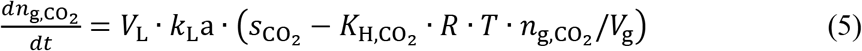

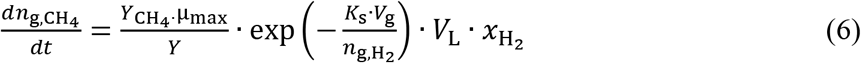

where *s*_CO_2__ is the concentration (mol/L) of carbon dioxide in the liquid phase and *x*_H_2__ is the biomass concentration (mol/L) of hydrogenotrophic methanogens. The numbers of moles in the gas phase are represented by the variables *n*_g,H_2__, *n*_g,CO_2__, *n*_g,CH_4__. The gas phase volume *V*_g_ = 20 mL and the liquid phase volume *V*_L_ = 6 mL. Liquid-gas transfer for carbon dioxide is described by a non-equilibria transfer rate which is driven by the gradient of the concentration of the gases in the liquid and gas phase. The transfer rate is determined by the mass transfer coefficient *k*_L_a (h^-1^) and the Henry’s law coefficients *K*_H,CO_2__ (M/bar). *R* (bar·(M · K)^-1^) is the ideal gas law constant and *T* is the temperature (K). Microbial decay is represented by a first-order kinetic rate with *k*_d_ (h^-1^) the death cell rate constant. Microbial growth was represented by the rate function proposed by Desmond-Le Quéméner and Bouchez (33) using hydrogen as the limiting reactant

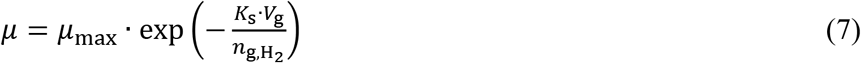

where *μ* is the growth rate (h^-1^), μ_max_ (h^-1^) is the maximum specific growth rate constant and *K*_s_(mol/L) the affinity constant. Equation (7) is derived from energetic principles following Boltzmann statistics and uses the concept of exergy (maximum work available for a microorganism during a chemical transformation). The affinity constant has an energetic interpretation since it is defined as

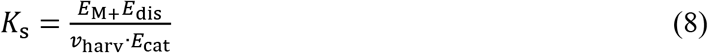

where *E*_dis_ (kJ/mol) and *E*_M_ (kJ/mol) are the dissipated exergy and stored exergy during growth respectively. *E*_cat_ (kJ/mol) is the catabolic exergy of one molecule of energy-limiting substrate, and *ν*_harv_ is the volume at which the microbe can harvest the chemical energy in the form of substrate molecules (33). *E*_cat_ is the absolute value of the Gibbs energy of catabolism (ΔG_r,c_) when the reaction is exergonic (ΔG_r,c_<0) or zero otherwise. The stored exergy *E*_M_ is calculated from a reaction (destock) representing the situation where the microbe gets the energy by consuming its own biomass. *E*_M_ is the absolute value of the Gibbs energy of biomass consuming reaction (ΔG_r,destock_) when the reaction is exergonic (ΔG_r,destock_<0) or zero otherwise. Finally, the dissipated exergy *E*_dis_ is the opposite of the Gibbs energy of the overall metabolic reaction, which is a linear combination of the catabolic and destock reactions. This calculation follows the Gibbs energy dissipation detailed in Kleerebezem and Van Loosdrecht (31).

In our model, the stoichiometry of methanogenesis is represented macroscopically by one catabolic reaction (R_1_) for methane production and one anabolic reaction (R_2_) for microbial formation. We assumed that ammonia is the only nitrogen source for microbial formation. The molecular formula of microbial biomass was assumed to be C_5_H_7_O_2_N (41).

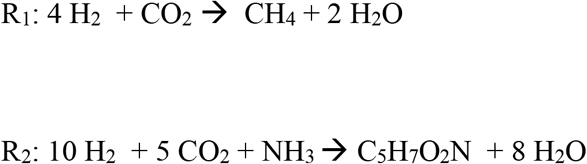

In the model, the stoichiometry of the reactions is taken into account *via* the parameters *Y*, *Y*_CO_2__, *Y*_CH_4__, which are the yield factors (mol/mol) of microbial biomass, CO_2_ and CH_4_. The microbial yield factor *Y* was extracted from literature. We assumed that *M. smithii* and *M. ruminantium* have the same yield (being both Methanobrevibacter). This yield factor was set to 0.006 mol biomass/mol H_2_, using the methane-based molar growth yield of 2.8 g biomass/mol CH_4_ estimated for *M. smithii* (42) and the equations (9) and (11). Similarly, the yield factor for *M. formicium* was set to 0.007 mol biomass/mol H_2_ using the methane-based molar growth yield of 3.5 g biomass/mol CH_4_ reported by Schauer and Ferry (43). The fraction of H_2_ utilized for microbial growth (reaction R_2_) is defined by the yield factor *Y* (mol of microbial biomass/mol of H_2_). Now, let *f* be the fraction of H_2_ used for the catabolic reaction R_1_. Reaction R_2_ tells us that for every 10 moles of H_2_ used in R_2_, we get 1 mol of microbial biomass. Hence, it follows that

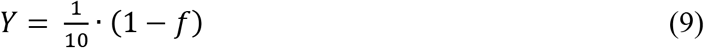

If *Y* is known, the fraction *f* can be computed from Equation (9).

The yield factors of CO_2_ and CH_4_ can be expressed as functions of the the fraction *f*:

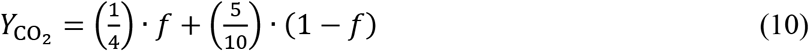

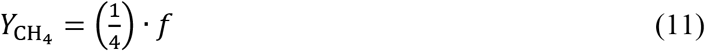

The model has two physicochemical parameters (*k*_L_a, *K*_H,CO_2__) and four biological parameters (*μ*_max_, *K*_s_, *Y*, *k*_d_). The initial condition for *s*_CO_2__ is unknown and was also included in the parameter vector for estimation. The Henry’s law coefficients are known values calculated at 39°C using the equations provided by Batstone et al. (41). If the article is accepted, an implementation of the model in the Open Source software Scilab (https://www.scilab.org/) will be made available at Zenodo (https://doi.org/10.5281/zenodo.3271611).

#### Theoretical model to study interactions among methanogens

The experimental study of microbial interactions requires sophisticated *in vitro* systems under continuous operation such as the one developed by Haydock *et al*. (44). In our work, we explored by means of mathematical modelling how the methanogens can interact under *in vivo* conditions. For this theoretical study, we elaborated a toy model based on the previous model for *in vitro* methanogenesis. Let us consider the following simple model for representing the consumption of hydrogen by the methanogenic species *i* under an *in vivo* scenario of continuous flow

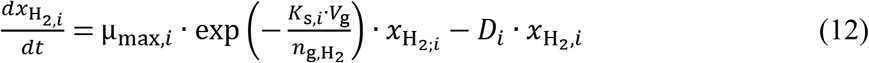

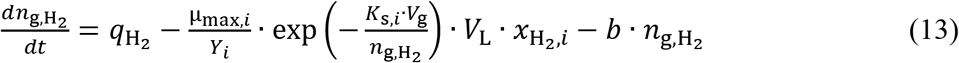

where *q*_H_2__ (mol/h) is the flux of hydrogen produced from the fermentation of carbohydrates. The kinetic parameters are specific to the species *i* (*x*_H_2,*i*__.). The parameter *D_i_* (h^-1^) is the dilution rate of the methanogens and *b* (h^-1^) is an output substrate rate constant. Extending the model to *n* species with a common yield factor *Y*, the dynamics of hydrogen is given by

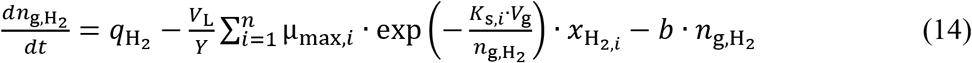

where the sub index *i* indicates the species. In our case study, *n* = 3.

#### Parameter identification

Before tackling the numerical estimation of the model parameters, we addressed the question of whether it was theoretically possible to determine uniquely the model parameters given the available measurements from the experimental setup. This question is referred to as structural identifiability (45). Structural identifiability analysis is of particular relevance for model whose parameters are biologically meaningful, since knowing the actual value of the parameter is useful for providing biological insight on the system under study (46). Moreover, in our case, we are interested in finding accurate estimates that can be further used as priors in an extended model describing the *in vivo* system.

We used the freely available software DAISY (47) to assess the structural identifiability of our model. Physical parameters (*k*_L_a, *K*_H,CO_2__) were set to be known. The model was found to be structurally globally identifiable. In practice, however, to facilitate the actual identification of parameters and reduce practical identifiability problems such as high correlation between the parameters (48), we fixed some model parameters to values reported in the literature. The transport coefficient *k*_L_a, the Henry’s law coefficient *K*_H,CO_2__, and the dead cell rate constant *k*_d_ were set to be known and were extracted from Batstone et al. (41). Therefore, only the parameters *μ*_max_, *K*_s_ and initial condition of *s*_CO_2__ were set to be estimated. To capitalize on the calorimetric data, we further assumed that *μ*_max_ was equal to the specific rate constant μ_c_ estimated from the heat flux-time curve. By this, only the affinity constant for each strain and the initial condition of *s*_CO_2__ were left to be estimated.

The parameter identification for each methanogen was performed with the IDEAS Matlab^®^ toolbox (49), which is freely available at http://genome.jouy.inra.fr/logiciels/IDEAS. The measured variables are the number of moles in the gas phase (H_2_, CH_4_, CO_2_). The Lin’s concordance correlation coefficient (CCC) (50) was computed to quantify the agreement between the observations and model predictions.

## Results

### Methanogens biomass

Archaea-specific primers targeting the 16S rRNA gene were used to enumerate microbial cells in each pure culture. Three hours post inoculation microbial numbers varied from 7.62×10^7^ to 2.81×10^8^ and reached 10^9^ after 72 hours of incubation. Table S3 summarizes microbial numbers at different sampling times.

### Calorimetric pattern of methanogens

Figure 2 displays a representative isothermal calorimetric curve for each methanogen. The three measured heat flux dynamics of each methanogen were found to follow similar energetic patterns. *M. smithii* and *M. formicium* exhibited a lag phase of a few hours, while *M. ruminantium* was already metabolically active when introduced into the minicalorimeter though several attempts were made to obtain a lag phase by changing storage conditions and thawing the culture just before inoculating the microcalorimetric ampoules. The pattern of heat flux for all tested methanogens is characterized by one predominant peak which was observed at different times for each methanogen. *M. smithii* exhibited a second metabolic event occurring at 60 h with an increase of heat flux. The same phenomenon was observed for *M. formicium* but at a lower intensity that started at 140 h. One possible explanation for this event is cell lysis (39). The process was considered completed when the heat flux ceased marking the end of the metabolic activity. It is noted that *M. formicium* produced a small peak at 14 h (Fig. 2). A similar peak, but of much smaller size, was observed on the other curves obtained with this methanogen. *M. smithii* also exhibits a small peak (occurrence of 3 out of 3) at 7.4 h shown in the inset of Figure 2. For *M. ruminantium*, we do not know whether the small peak exists since the initial part of the curve is missing. This small peak translates in a metabolic activity that remains to be elucidated.

**Figure 2.**
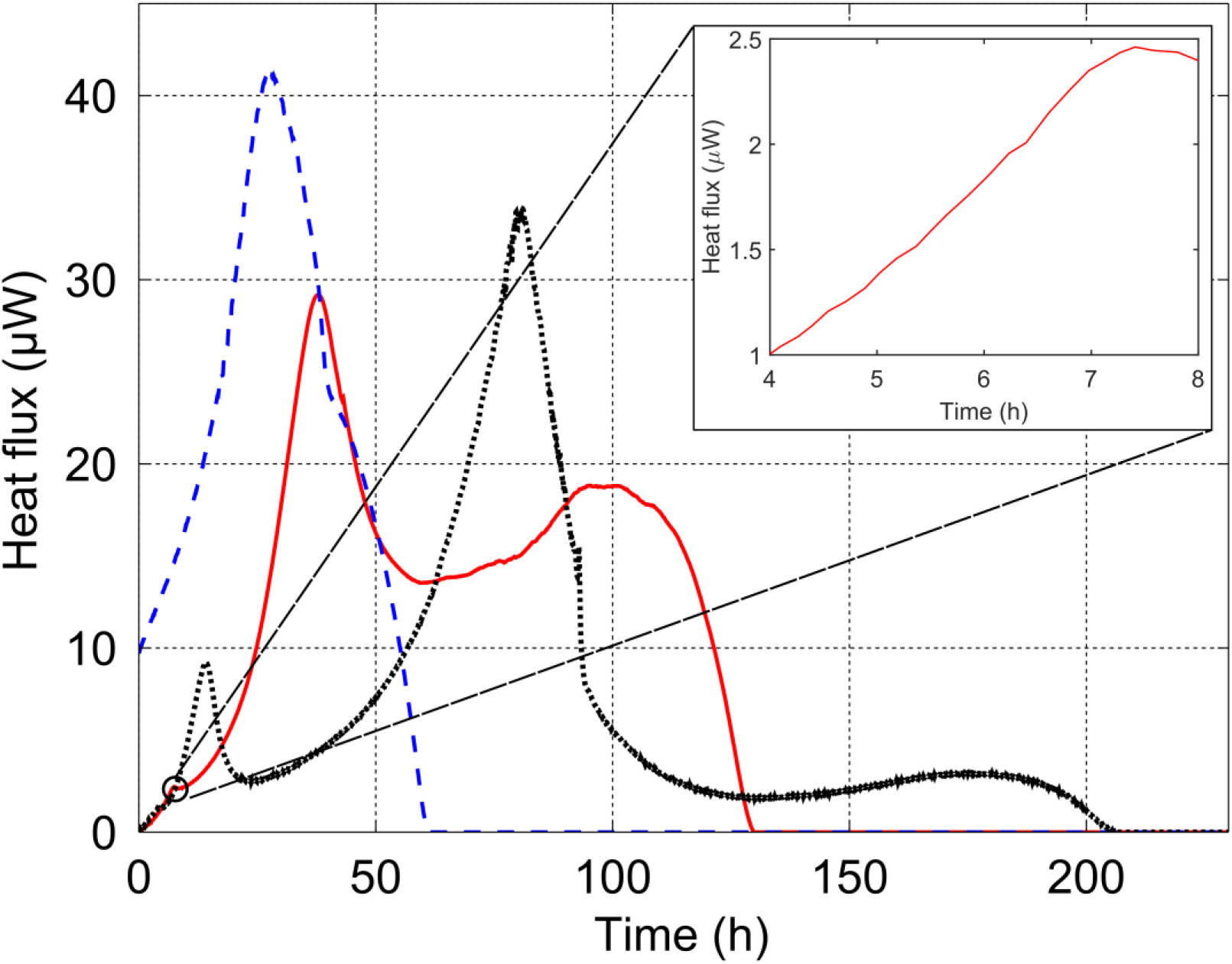
Example of isothermal calorimetric curves for *M. ruminantium* (dashed blue line), *M. smithii* (solid red line) and *M. formicium* (dotted black line). The dominant metabolic phase is represented by one peak. The magnitude of the peak differs between the methanogens and also the slope of the heat flux trajectories. The return of the heat flux to the zero baseline also differs between the three methanogens. The inset zoom displays the peak exhibited by *M. smithii* at 7.4 h.

The total heat (*Q_m_*) produced during the methanogenesis process that took place under the present experimental conditions was, on average, −5.5 ± 0.5 J for the three methanogens (for *M. ruminantium*, the missing initial part of the heat flux-time curve was approximately estimated by extrapolating the exponential fit). As we shall see below, this experimental value is consistent with the theoretically expected value.

### Estimation of thermodynamic properties

We defined two macroscopic reactions to represent the catabolism (R1) and anabolism (R2) of the methanogenesis (see Modelling in vitro methanogenesis section). All thermodynamic properties result from the contribution of both catabolic and anabolic reactions. In the Supplementary Material, the calculations of total heat (*Q*_m_), enthalpy (Δ*H*_m_), Gibbs energy (Δ*G*_m_) and entropy (Δ*S*_m_) of the methanogenesis are detailed. The estimated overall heat produced during the methanogenesis process under our experimental conditions was in average *Q*_m_ = −5.66 J. This heat results from the sum of the heat of the catabolic reaction (*Q*_c_) and the heat of the anabolic reaction (*Q*_a_). From the total heat of the methanogenesis, the anabolic reaction contributes to 7% of the metabolic heat for*M. smithii* and *M. ruminantium*. For *M. formicium*, the contribution of the anabolic reaction to the metabolic heat is 9%. It is also interesting to note that there is a very good agreement between the theoretical value calculated above and the overall heat experimentally determined by microcalorimetry (−5.5 ± 0.5 J).

Table 1 shows the thermodynamic properties per mole of biomass formed during methanogenesis of *M. ruminantium, M. smithii* and *M. formicium* on H_2_/CO_2_. These properties are compared with values found in the literature for other methanogens grown on different substrates.

**Table 1.** Gibbs energies, enthalpies and entropies of metabolic processes involving some methanogens growing on different energy sources

| Microorganism | Energy<br>substrate | Growth<br>conditions | $\Delta G_m$<br>kJ / C-mol | $\Delta H_m$<br>kJ / C-mol | $T\Delta S_m$<br>kJ / K<sup>-1</sup> C-mol | Driving force | Reference |
| --- | --- | --- | --- | --- | --- | --- | --- |
| *M. ruminantium*,<br>*M. smithii*, | H<sub>2</sub>/CO<sub>2</sub> | anaerobic | -1073 | -2132 | -1059 | Enthalpy-driven<br>but Entropy-retarded | this<br>work |
| *M. formicium* | H<sub>2</sub>/CO<sub>2</sub> | anaerobic | -801 | -1605 | -804 | Enthalpy-driven<br>but Entropy-retarded | this<br>work |
| *M. thermo-*<br>*autotrophicum* | H<sub>2</sub>/CO<sub>2</sub> | anaerobic | -802 | -3730 | -2928 | Enthalpy-driven<br>but Entropy-retarded | (53) |
| *M. formicium* | formate | anaerobic | -880 | -613 | +267 | Enthalpy-driven | (30) |
| *M. barkeri* | methanol | anaerobic | -570 | -420 | +150 | Enthalpy-driven | (30) |
| *M. barkeri* | acetate | anaerobic | -366 | +145 | +511 | Entropy-driven<br>but enthalpy-retarded | (76) |

### Dynamic description of *in vitro* kinetics

The developed mathematical model was calibrated with the experimental data from *in vitro* growth experiments in Balch tubes. Table 2 shows the parameters of the dynamic kinetic model described in Equations 2-6. The reported value of *μ*_max_ for each methanogen corresponds to the average value obtained from three heat flux-time curves. From Table 2, it is concluded that *M. smithii* exhibited the highest growth rate constant, followed by *M. ruminantium* and finally *M. formicium*. In terms of the affinity constant *K*_s_, while *M. smithii* and *M. ruminantium* are of the same order, the affinity constant for *M. formicium* is lower in one order of magnitude.

**Table 2.** Parameters of the model of *in vitro* methanogenesis. The value reported *μ*_max_ for each methanogen is the mean value obtained from three heat flux-time curves

| Parameter | Definition | Value |  |  |
| --- | --- | --- | --- | --- |
| $k_L a$ (h<sup>-1</sup>) | Liquid–gas transfer constant | 8.33 | | |
| $K_{H,CO_2}$ (M/bar) | Henry’s law coefficient of carbon dioxide | 0.0246 | | |
| $k_d$ (h<sup>-1</sup>) | Death cell rate constant | $8.33 \times 10^{-4}$ | | |
|  |  | *M. smithii* | *M. ruminantium* | *M. formicium* |
| $K_s$ (mol/L) | Affinity constant | 0.028 | 0.042 | 0.011 |
| $\mu_{\max}$ (h<sup>-1</sup>) | Maximum specific growth rate constant | 0.12 | 0.07 | 0.046 |
| $Y$ (mol biomass /mol H<sub>2</sub>) | Microbial biomass yield factor | 0.006 | 0.006 | 0.007 |

Figure 3 displays the dynamics of the compounds in the methanogenesis for the three methanogens. Experimental data are compared against the model responses. Table 3 shows standard statistics for model evaluation. The model captures efficiently the overall dynamics of the methanogenesis. Hydrogen and methane are very well described by the model with concordance correlation coefficients (CCC) of 0.99 and 0.97 respectively. For carbon dioxide, CCC = 0.94.

**Figure 3.**
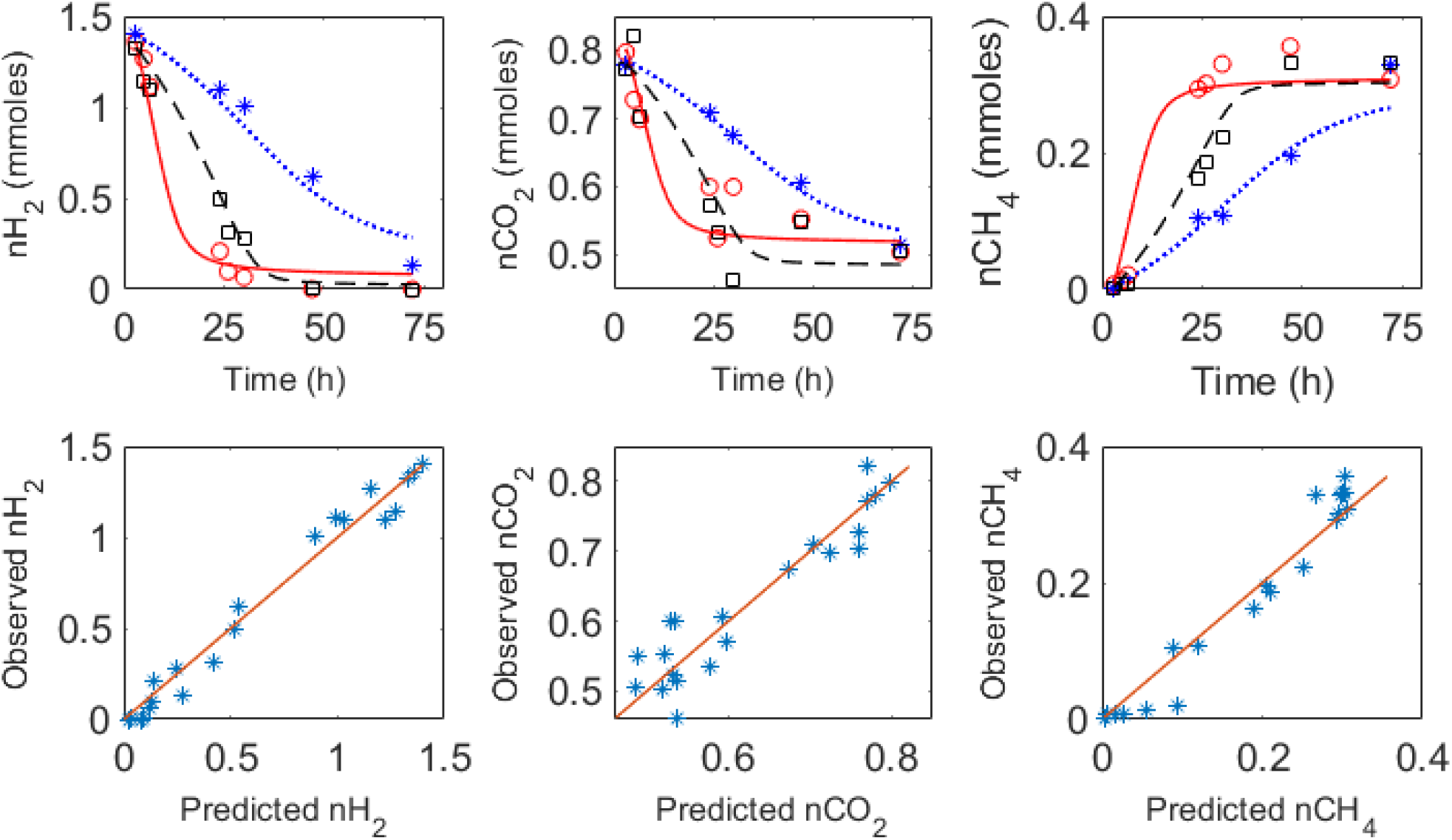
Top plots: dynamics of methanogenesis by *M. ruminantium* (*), *M. smithii* (o) and *M. formicium* (□). Experimental data (*,o,□) are compared against model predicted responses: dotted blue lines (*M. ruminantium*), solid red lines (*M. smithii*) and dashed black lines *(M. formicium)*. Bottom plots: summary observed vs predicted variables. The solid red line is the isocline.

**Table 3.** Statistical indicators for model evaluation

| Component | CCC* | $r^2$ | CV<sub>RMSE</sub>** |
| --- | --- | --- | --- |
| Hydrogen | 0.99 | 0.97 | 14 |
| Methane | 0.97 | 0.95 | 18 |
| Carbon dioxide | 0.94 | 0.88 | 6 |

\* CCC: Lin's concordance correlation coefficient.
\*\* CV<sub>RMSE</sub>: coefficient of variation of the root mean squared error.

Figure 4 displays the dynamics for the methanogens as measured by the 16S rRNA gene, as well as the dynamics of biomass as predicted for the model. As observed, the microbes follow a typical Monod-like trajectory.

**Figure 4.**
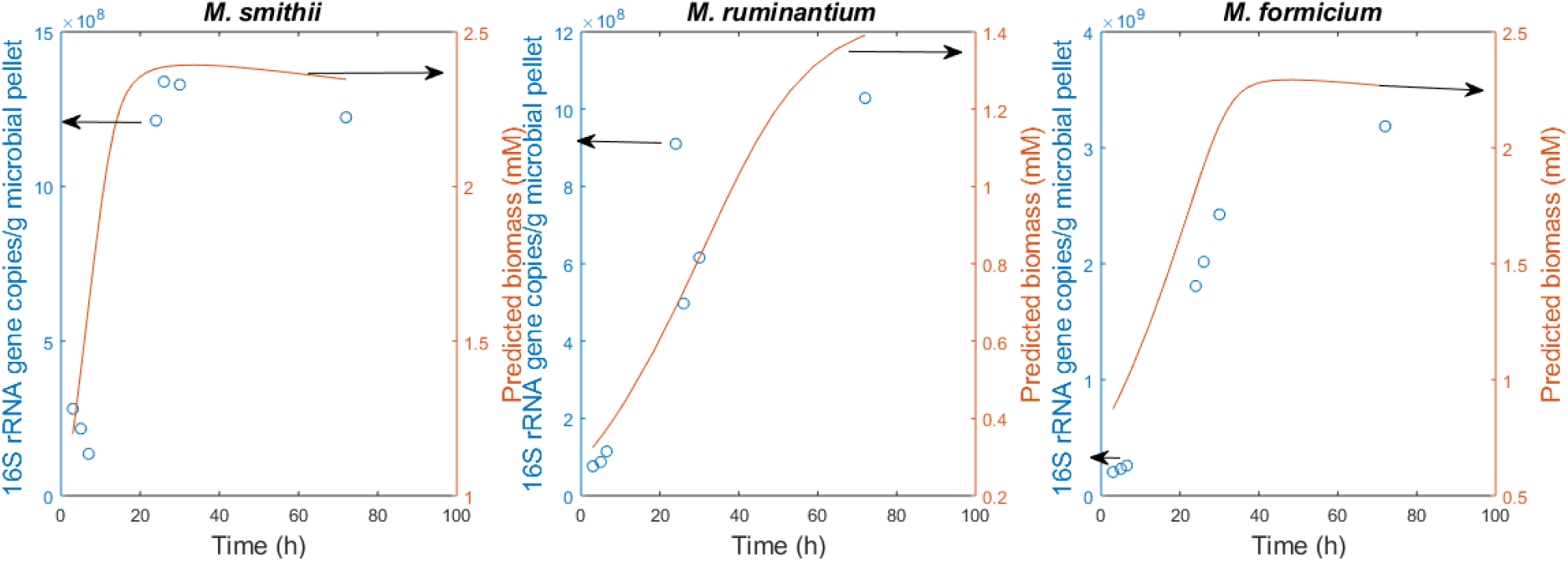
Dynamics of methanogens as measured by 16S rRNA gene copies (circles) and biomass concentrations (solid line) predicted by the model.

## Discussion

Our objective in this work was to quantitatively characterize the dynamics of hydrogen utilization, methane production, growth and heat flux of three hydrogenotrophic methanogens by integrating microbiology, thermodynamics and mathematical modelling. Our model developments were instrumental to quantify energetic and kinetic differences between the three methanogens studied, strengthening the potentiality of microcalorimetry as a tool for characterizing the metabolism of microorganisms (51). This modelling work provides estimated parameters that can be used as prior values for other modelling developments of gut microbiota.

### Energetic and kinetic differences between methanogens

Methanogenesis appears as a simple reaction with a single limiting substrate (H_2_). The microcalorimetry approach we applied revealed that this simplicity is only apparent and that hydrogenotrophic methanogens exhibit energetic and kinetic differences. Methanogenesis is indeed a complex process that can be broken down in several stages. The dominant metabolic phase is represented by one peak that occurs at different times. The magnitude of the peak differs between the methanogens and also the slope of the heat flux trajectories. The return time of the heat flux to the zero baseline was also different. The energetic difference is associated with kinetic differences that translate into specific kinetic parameters, namely affinity constant (*K*_s_) maximum growth rate constant (μ_max_). Previously, energetic differences between methanogens have been ascribed to the presence or absence of cytochromes (14). These differences are translated into different yield factors, H_2_ thresholds, and doubling times. The kinetic differences revealed in this study for three cytochrome-lacking methanogens indicate that factors other than the presence of cytochromes might play a role in the energetics of methanogenesis. Interestingly, calorimetric experiments showed that *M. ruminantium* was metabolically active faster than the other methanogens, characteristic that could explain the predominance of *M. ruminantium* in the rumen (52). Looking at the expression of the affinity constant (Equation (8)), the differences between the affinity constants among the methanogens can be explained by the differences between the by the harvest volume *ν*_harv_ and the yield factors. Note that in the kinetic function developed by Desmond-Le Quéméner and Bouchez (33), the maximum growth rate did not have any dependency on the energetics of the reaction. Our experimental study revealed that *μ*_max_ is species-specific and reflects the dynamics of the heat flux of the reaction at the exponential phase. This finding suggests that a further extension of the kinetic model developed by Desmond-Le Quéméner and Bouchez (33) should include the impact of energetics on *μ*_max_. Since our study is limited to three species, it is important to conduct further research on other methanogens to validate our findings.

### Thermodynamic analysis

Regarding the energetic information for different methanogens summarized in Table 1, it is observed that the thermodynamic behaviour of the three methanogens is analogous to that observed for *Methanobacterium thermoautotrophicum* (53). The values reported in Table 1 show indeed that the methanogenesis on H_2_/CO_2_ is characterized by large heat production. The growth is highly exothermic, with a *ΔH_m_* value that largely exceeds the values found when other energy substrates are used. The enthalpy change *ΔH_m_*, which is more negative than the Gibbs energy change *ΔG_m_*, largely controls the process. Growth on H_2_/CO_2_ is also characterized by a negative entropic contribution *TΔS_m_* which, at first sight, may look surprising since entropy increases in most cases of anaerobic growth (54). However, this can be understood if one remembers that *TΔS_m_* corresponds in fact to the balance between the final state and the initial state of the process, that is

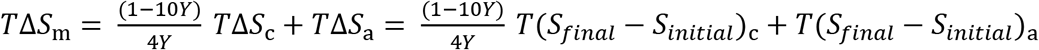

Methanogenesis on H_2_/CO_2_ is particular because the final state of its catabolic reaction (1 mol CH_4_ + 2 mol H_2_O) involves a smaller number of moles than the initial state (4 mol H_2_ + 1 mol CO_2_), which results in a significant loss of entropy during the process. For spontaneous growth in such a case, the Δ*H*_m_ must not only contribute to the driving force but must also compensate the growth-unfavourable TΔ*S*_m_, which means that Δ*H*_m_ must be much more negative than Δ*G*_m_ (55). For this reason, methanogenesis on H_2_/CO_2_, which is accompanied by a considerable decrease of entropy and a large production of heat, has been designed as an entropy-retarded process (53). More generally, von Stockar and Liu (55) noticed that when the Gibbs energy of the metabolic process is resolved into its enthalpic and entropic contributions, very different thermodynamic behaviours are observed depending on the growth type. These thermodynamic behaviours are: aerobic respiration is clearly enthalpy-driven (Δ*H*_m_ << 0 and TΔ*S*m > 0), whereas fermentative metabolism is mainly entropy-driven (Δ*H*_m_ < 0 and TΔ*S*_m_ >> 0). Methanogenesis on H_2_/CO_2_ is enthalpy-driven but entropy-retarded (Δ*H*_m_ << 0 and TΔ*S*m < 0), whereas methanogenesis on acetate is entropy-driven but enthalpy-retarded (Δ*H*_m_ > 0 and TΔ*S*_m_ >> 0). In the present case, the highly exothermic growth of *M. ruminantium, M. smithii* and *M. formicium* on H_2_/CO_2_ is largely due to the considerable decrease of entropy during the process: in fact, 50% of the heat produced here serves only to compensate the loss of entropy. A proportion of 80% was found for *M. thermoautotrophicum* (53), which results from the fact that their TΔ*S*_m_ and Δ*H*_m_ values are, respectively, 2.7 and 1.7 times larger than ours. This difference might be due to the differences in the temperature of the studies, namely 39°C in our study vs 60°C in the study by Schill et al. (53).

### Do our results inform on ecological questions such as species coexistence?

The competitive exclusion principle (56) states that coexistence cannot occur between species that occupy the same niche (the same function). Only the most competitive species will survive. Recently, by using thermodynamic principles, Großkopf & Soyer (34) demonstrated theoretically that species utilizing the same substrate and producing different compounds can coexist by the action of thermodynamic driving forces. Since in our study the three methanogens perform the same metabolic reactions, the thermodynamic framework developed Großkopf & Soyer (34) predicts, as the original exclusion principle (56), the survival of only one species. By incorporating thermodynamic control on microbial growth kinetics, Lynch et al (57) showed theoretically that differentiation of ATP yields can explain ecological differentiation of methanogens over a range of liquid turnover rates. This theoretical work predicts that for a fixed liquid turnover rate, only one species survives. For the continuous culture of microorganisms, it has been demonstrated that at the equilibrium (growth rate equals the dilution rate) with constant dilution rates and substrate input rates, the species that has the lowest limiting substrate concentration wins the competition. From Eq. (12), the number of moles of hydrogen of the species 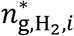 at the steady state is

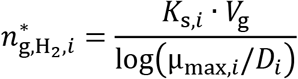

Using the model parameters of Table 2, we studied *in silico* three possible competition scenarios, assuming a constant environment (constant dilution rate *D*). Two dilution rates were evaluated: *D* = 0.021 h^-1^ (retention time = 48 h) and *D* = 0.04 h^-1^ (retention time = 25 h). A retention time of 48 h corresponds to values measured in small ruminants (58) and to humans as we used in our gut model (20). For higher retention times, the results obtained for 48 h hold. For *D* = 0.021 h^-1^, we obtained that 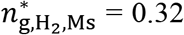 mmol, 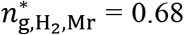 mmol, 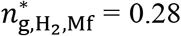 mmol, where the subindex Ms, Mr, Mf stand for *M. smithii, M. ruminantium* and *M.formicium*. From these results, it appears that under a constant environment, *M. formicium* will win the competition. Since 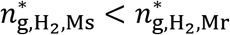, *M. ruminantium* will be extinguished before *M. smithii*. For *D* = 0.04 h^-1^, we obtained that 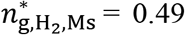 mmol, 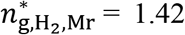 mmol, 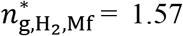 mmol, and thus *M. smithii* wins the competition. To win the competition, *M. ruminantium* requires longer retention times than its competitors. Retention times of digesta longer than 48 h are physiologically uncommon, thus the presence of *M. ruminantium* in the gut ecosystem can be explained, for example, from known adhesion properties (both *M. ruminantium* and *M. smithii* genes encode adhesin-like proteins (59,60). To illustrate these aspects, we built a multiple-species model with the three methanogens using Eq. (12) and Eq. (14). The parameter *b* was set to 0.5 h^-1^ and the hydrogen flux production *q*_H_2__ rate was set to 0.02 mol/min. Figure 5A displays the dynamics of the three methanogens for the first scenario (*D*= 0.021 h^-1^). It is observed that at 50 d only *M. formicium* survives. This result, however, is not representative of what occurs in the rumen where the three methanogens coexist (5,61). It is intriguing that in our toy model it is *M. formicium* that wins the competition, bearing in mind that *M. ruminantium* and *M. smithii* are more abundant than *M. formicium* (5,52). Figure 5 shows that selective conditions favour the survival of one species. Similar results can be obtained for the human gut by including the effect of pH on microbial growth (23) and setting the gut pH to select one of the species. On the basis of the competitive exclusion principle, it is thus intriguing that having a very specialized function, methanogens are a diverse group that coexist. Gut ecosystems, therefore, exhibit the paradox of the plankton introduced by Hutchinson (1961) that presents the coexistence of species all competing for the same substrate in a relatively isotropic or unstructured environment (62). In the case of the rumen, our modelling work suggests that in addition to kinetic and thermodynamic factors, other forces contribute to the ecological shaping of the methanogens community in the rumen favouring the microbial diversity. Indeed, methanogenic diversity in the rumen results from multiple factors that include pH sensitivity, the association with rumen fractions (fluid and particulate material), and the endosymbiosis with rumen protozoa (5,52). For the human gut, ecological factors enable methanogens to coexist to a competitive environment where hydrogenotrophic microbes (acetogens, methanogenic archaea and sulfate-reducing bacteria) utilize H_2_ *via* different pathways (63–65). Both in the human gut and in the rumen, microbes grow in association with biofilms that form a polymer-based matrix that provides nutritional and hydraulic advantages for microbial growth and resistance to shear forces (20,66). Indeed, in our modelling work of human gut fermentation (20), we suggested that, from the different actions the mucus has on colonic fermentation, the mechanism of promoting conditions for microbial aggregation appears as the most relevant factor for attaining the high microbial density and the high level of fibre degradation characteristic of the human gut. Altogether, these factors result in nonlinear behaviours, spatial and temporal variations that promote coexistence and diversity, that, as discussed in dedicated literature on microbial ecology (67–71), render the classical formulation of the competitive exclusion principle (56,72) inapplicable to gut ecosystems.

**Figure 5.**
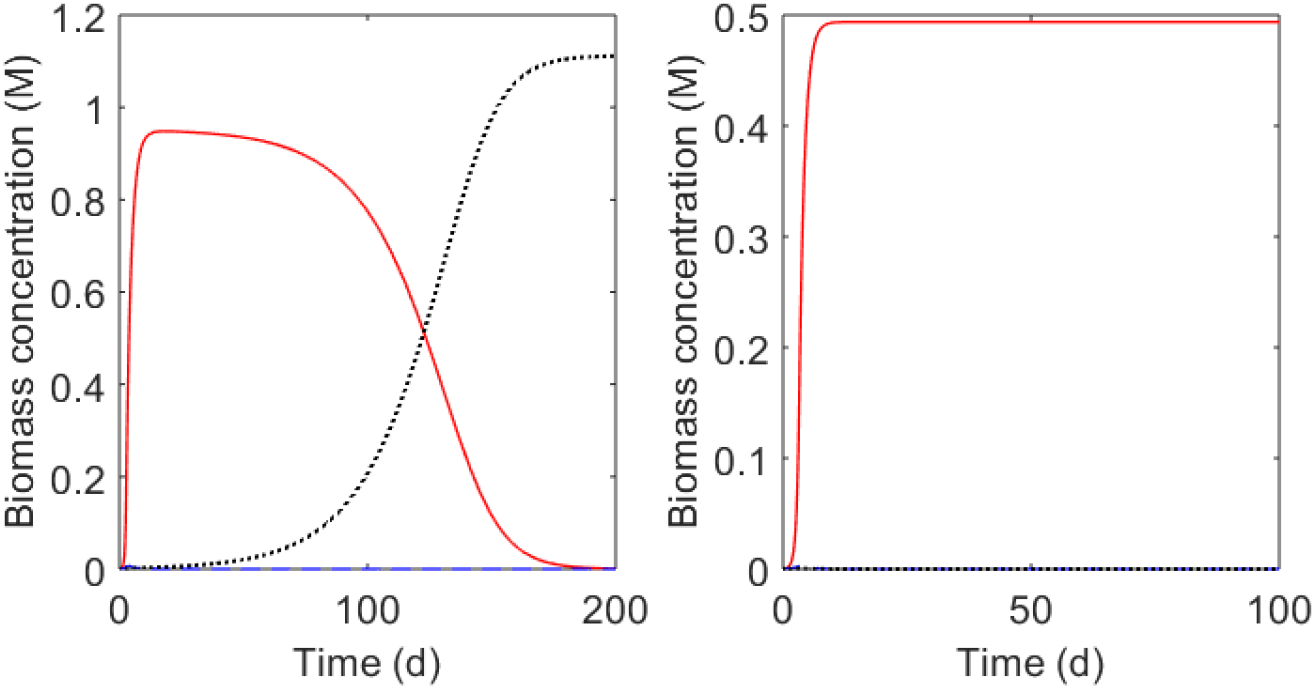
Possible competition scenarios between *M. ruminantium* (blue dashed line), *M. smithii* (red solid line) and *M. formicium* (black dotted line) in a hypothetical constant environment. A. At constant dilution rate of 0.021 h^-1^, *M. formicium* displaces the other two methanogens. B. With a constant dilution rate of 0.04 h^-1^, *M. smithii* wins the competition. At constant environmental conditions, only one species wins and displaces the other methanogens.

Finally, mathematical modelling is expected to enhance our understanding of gut ecosystems (66,73). It is then key that in addition to metabolic aspects, mathematical models of gut fermentation incorporate the multiple aspects that shape microbial dynamics to provide accurate predictions and improve insight on gut metabolism dynamics and its potential modulation. For ruminants, the development of precision livestock technologies provides promising alternatives for integrating real-time data of key animal phenotypes such as feeding behaviour with mathematical models for estimating methane emissions (74) and rumen function indicators at large scale. These tools will be instrumental to support livestock management decisions and guide timely interventions. Similarly, for humans, mathematical models coupled with electronic technologies for online monitoring of gut function (75) might facilitate the diagnosis and the design of personalized therapies for gastrointestinal diseases.

## Acknowledgements

We are grateful to Dominique Graviou (UMRH, Inra) for her skilled assistance on the *in vitro* growth experiments and qPCR assays. We thank the Inra PHASE department and the Inra MEM metaprogramme for financial support.

## Supplementary Material

### 1. Growth media information and inoculation conditions

**Table S1.**
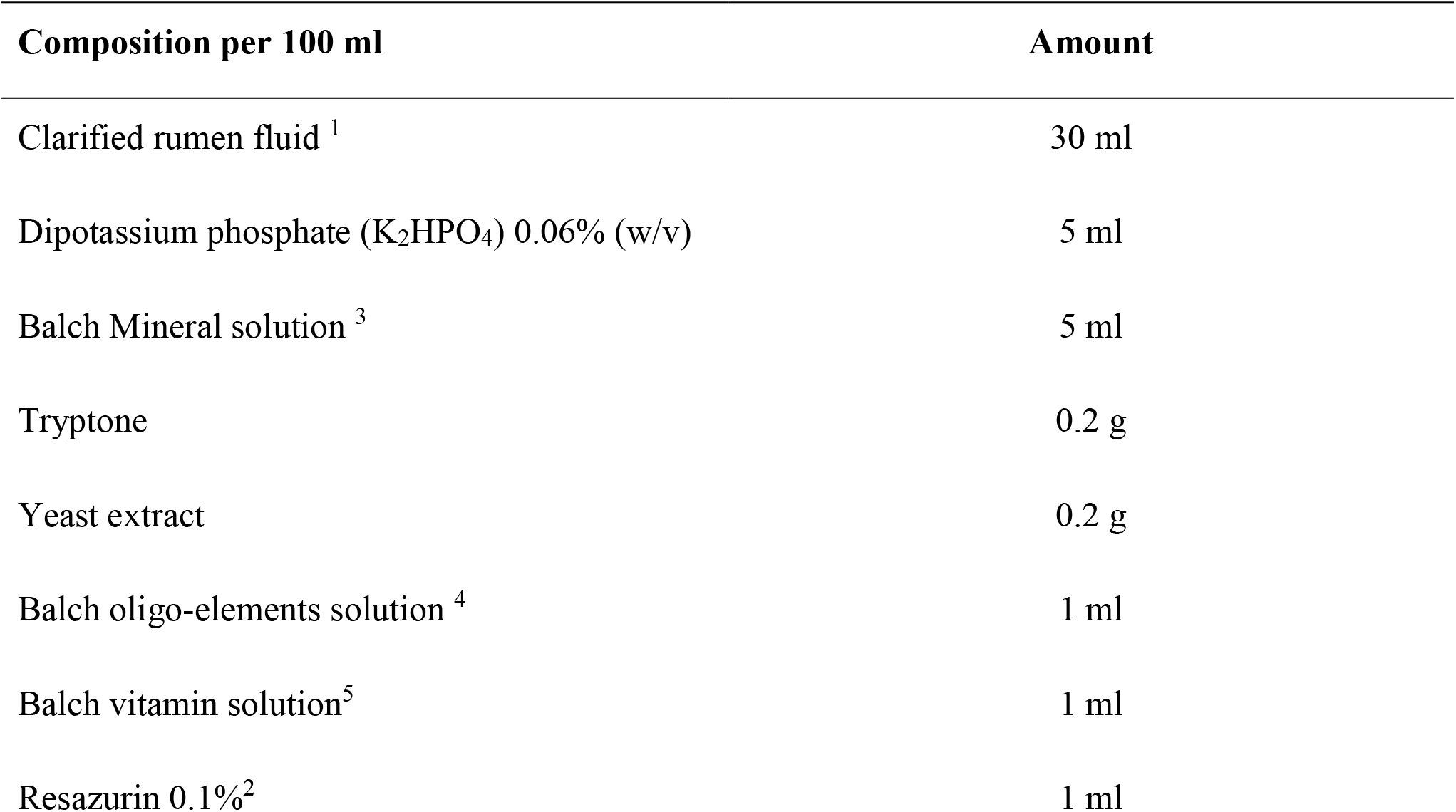
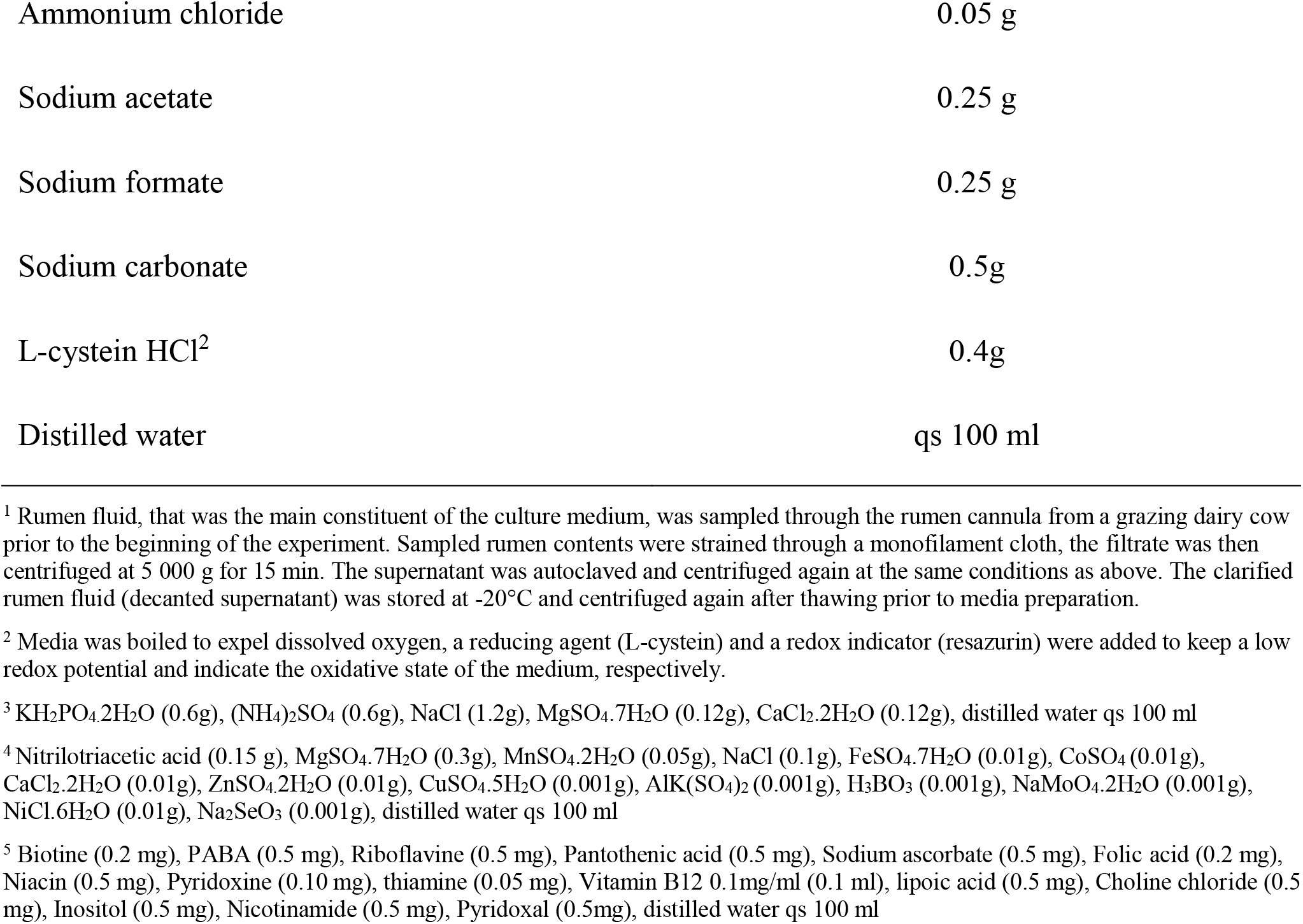
Methanogens growth media composition (modified from DSMZ medium 119 https://www.dsmz.de/microorganisms/medium/pdf/DSMZ_Medium119.pdf. Growth media was distributed in Balch tubes (6 ml per tube), tubes were sealed and sterilized by autoclaving at 121°C for 20 min. Media preparation and distribution was realized under CO_2_ flushing to assure anoxic conditions. Oxygen traces from commercial gases were scrubbed using a heated cylinder containing reduced copper (35).

**Table S2.**
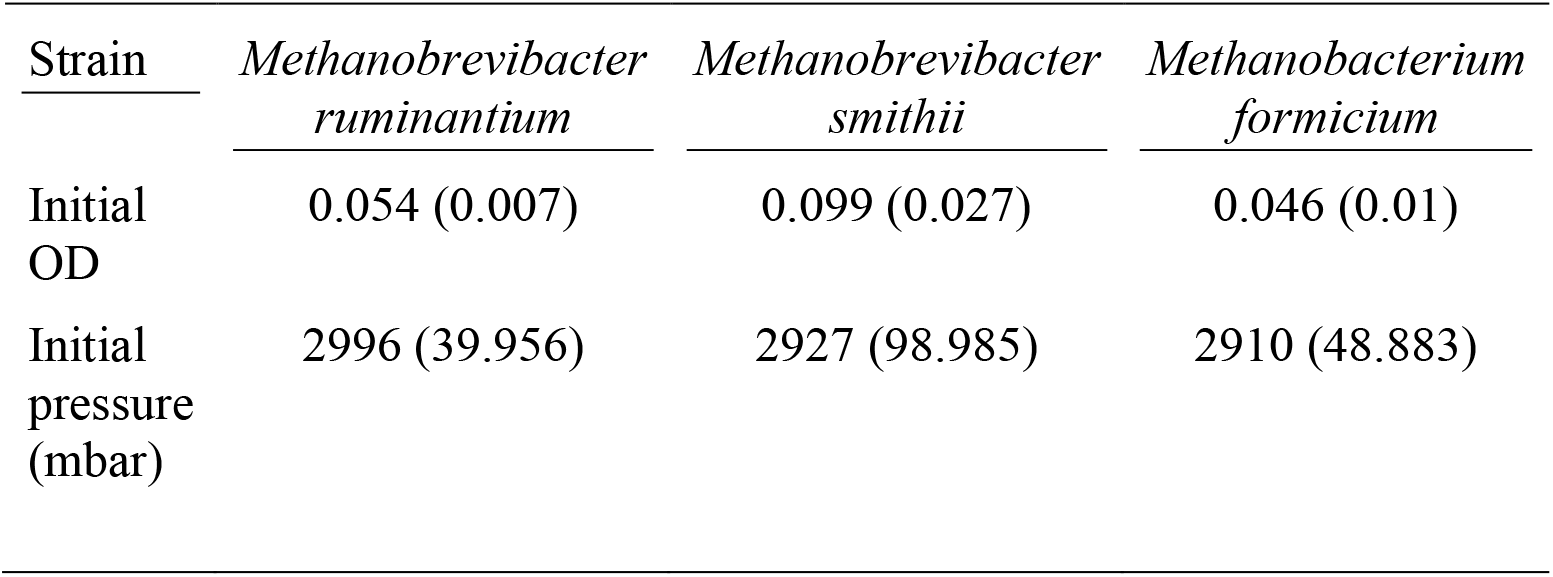
Summary of initial OD and pressure measured immediately after primary inoculation. Presented are means (sd) of 31 values per strain.

| Strain | *Methanobrevibacter ruminantium* | *Methanobrevibacter smithii* | *Methanobacterium formicium* |
| --- | --- | --- | --- |
| Initial OD | 0.054 (0.007) | 0.099 (0.027) | 0.046 (0.01) |
| Initial pressure (mbar) | 2996 (39.956) | 2927 (98.985) | 2910 (48.883) |

### 2. Quantification of methanogens

**Table S3.**
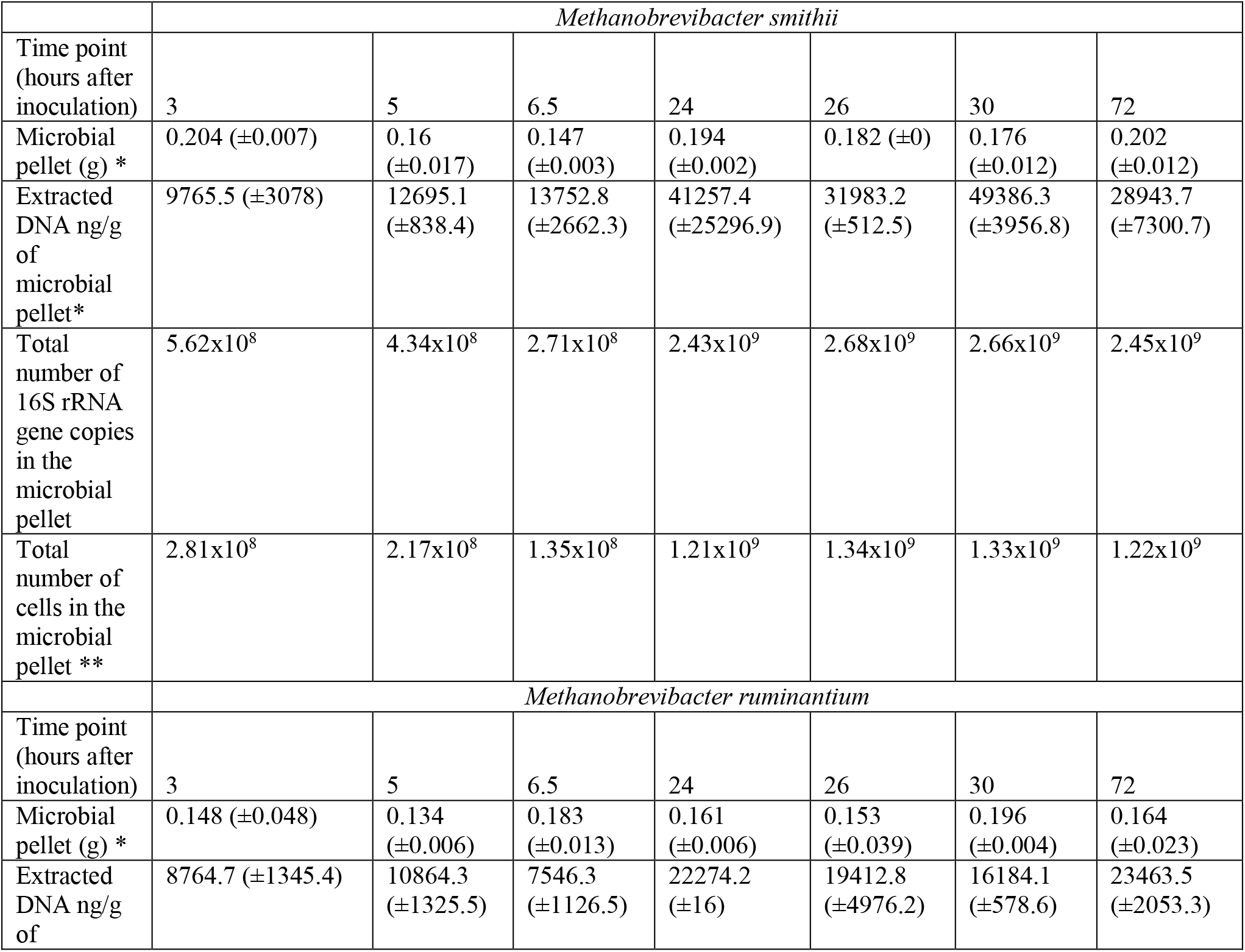
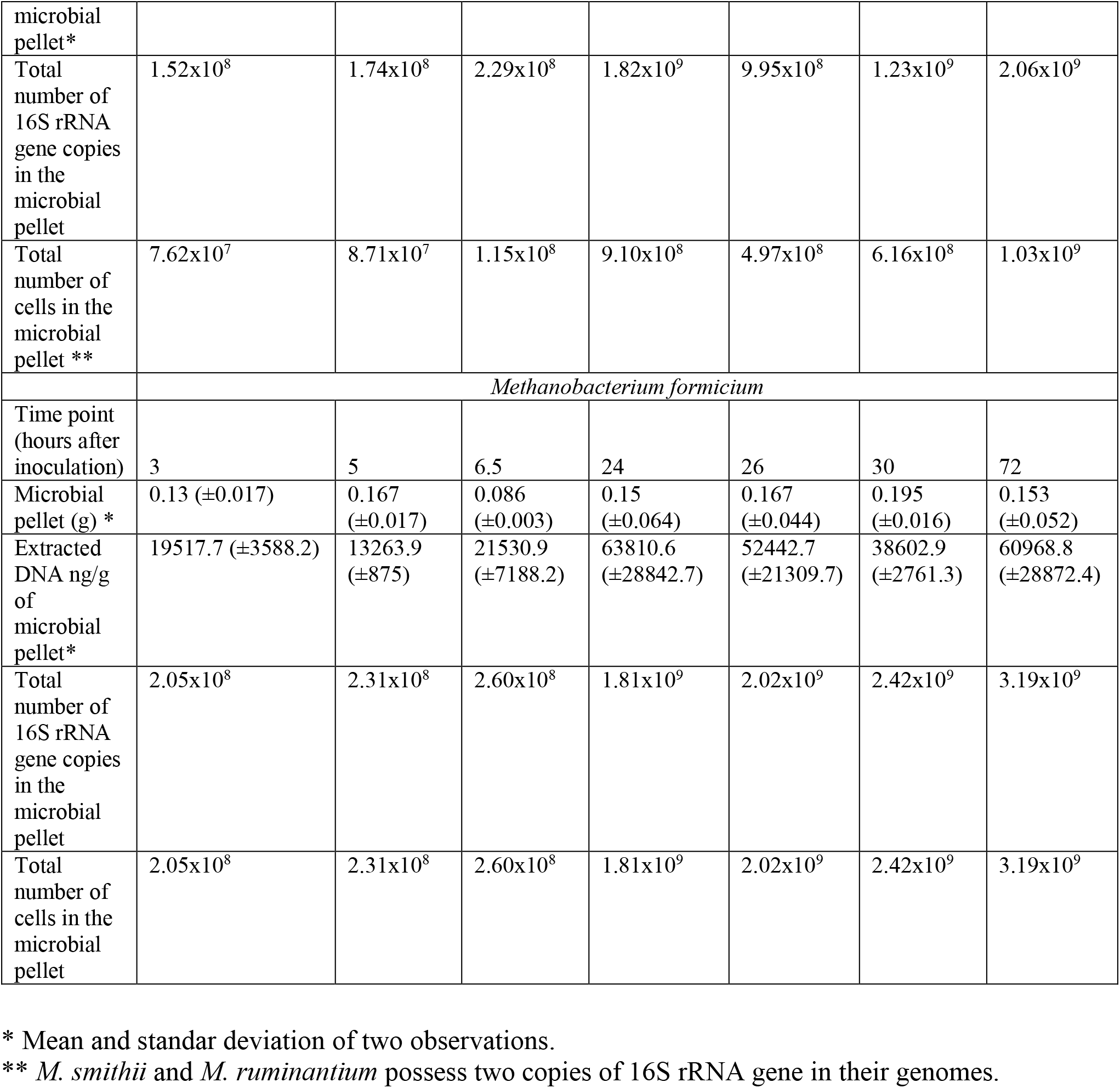
qPCR quantification of 16S rRNA genes

### 3. Calculation of thermodynamic properties of the methanogenesis

In our model, the methanogenesis is represented macroscopically by one catabolic reaction (R1) for methane production and one anabolic reaction (R2) for microbial formation. We assumed that ammonia is the only nitrogen source for microbial formation. The molecular formula of microbial biomass was assumed to be C_5_H_7_O_2_N (41).

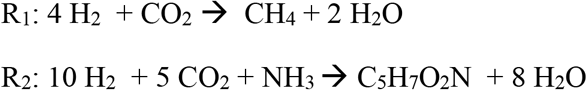

The thermodynamic properties associated to the methanogenesis result from the contribution of both catabolic and anabolic reactions.

#### 3.1 Thermodynamic properties of formation used for the calculations

**Table S4.** Standard enthalpies (Δ*H*f°) and Gibbs energies (Δ*G*f°) of formation at 25°C of compounds involved in hydrogenotrophic methanogenesis. Values were extracted from Wagman et al. (77), with the exception of the microbial biomass that was calculated from values for *Methanosarcina barkeri* reported by Liu et al. (76)

| Compound (phase) | $\Delta H_f^\circ$ (kJ/mol) | $\Delta G_f^\circ$ (kJ/mol) |
| --- | --- | --- |
| H<sub>2</sub>O (l) | -285.830 | -237.129 |
| H<sub>2</sub> (g) | 0 | 0 |
| CO<sub>2</sub> (g) | -393.509 | -394.359 |
| CH<sub>4</sub> (g) | -74.81 | -50.72 |
| NH<sub>3</sub> (aq) | -80.29 | -26.50 |
| C<sub>5</sub>H<sub>7</sub>O<sub>2</sub>N | -511.50* | -349.50* |

\* These values are five times the values reported by Liu et al. (76) since the biomass formula we used has five carbon molecules, while the biomass formula used by Liu et al. has one carbon.

#### 3.2 Enthalpies

The heat produced during methanogenesis results from the contribution of both catabolic and anabolic reactions. So, first, we calculated the standard enthalpies of the catabolic and anabolic reactions using the standard enthalpies of formation given in Table S4 for the different compounds involved in methanogenesis.

The standard enthalpy of the catabolic reaction 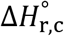 was calculated as follows

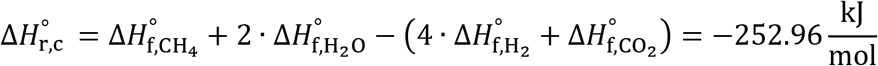

A similar equation was used for the calculation of the standard enthalpy of the anabolic reaction 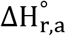

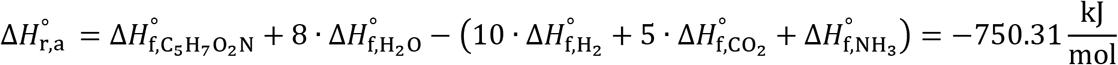

These results are at 25°C since this is the temperature of the standard enthalpies of formation reported in Table S3. A correction could be made to get results at 39°C but the heat capacities reported by Wagman et al. (77) show that the temperature correction can be neglected. Similarly, in the interest of simplicity, we assumed that the effect of pressure is negligible. Next, we considered the fact that the heat of a given reaction can be calculated at any state along the reaction pathway *via* the determination of the reaction coordinate or degree of advancement *ε* (78). Under our assumptions, the heat produced or consumed by a particular reaction during a given interval can be calculated as follows

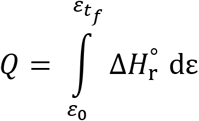

For our two reactions, at the instant *t* we have

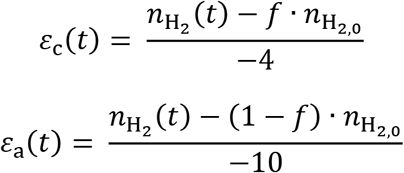

where *n*_H_2__ (*t*) is the number of moles of hydrogen at the instant *t*, *n*_H_2,0__ is the initial number of moles of hydrogen, and *f* is the fraction of H_2_ used for the catabolic reaction. Our calorimetric experiments started with *n*_H_2,0__ = 8.83 ≥ 10^-5^ mol in all cases. At the final time *t*_f_, all the hydrogen was consumed, so that *n*_H_2__(*t*_f_) = 0. For *M. smithii* and *M. ruminantium*, the microbial yield factor is *Y*=0.006 (6) which implies that *f* =0.94. Accordingly, *ε*_c_ = 2.075 · 10^-5^ mol and *ε*_a_ = 5.30 ≥ 10^-7^ mol. It thus follows that the overall heat produced during the methanogenesis process (*Q*_m_) can be calculated using the following equation

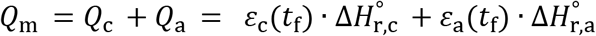

where *Q*_c_, *Q*_a_ are the heat produced during catabolism and anabolism respectively. The previous equation can also be written as

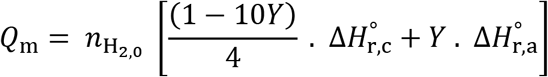

Under the experimental conditions of our study, this yields

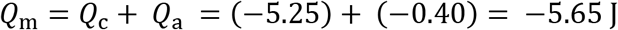

This result shows that the anabolic reaction contributes to only 7% of the metabolic heat.

Since the substrate was totally consumed, the enthalpy of the methanogenesis process per mole (or C-mol) of biomass formed, Δ*H*_m_, can be calculated as follows

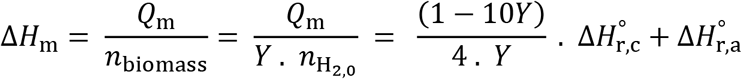

which yields

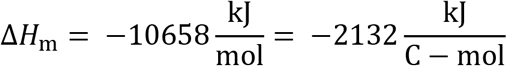

For*M. formicium, Y*=0.007 (7). Applying the same procedure, we obtained *Q*_m_ = −5.66 J and 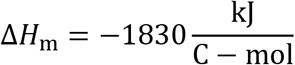 The anabolic reaction contributes to 8% of the metabolic heat.

#### 3.3 Gibbs energies and entropies

Following a procedure analogous to the one used above for the enthalpies, the standard Gibbs energies of the catabolic 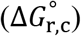 and anabolic 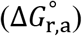 reactions were calculated using the standard Gibbs energies of formation listed in Table S3.

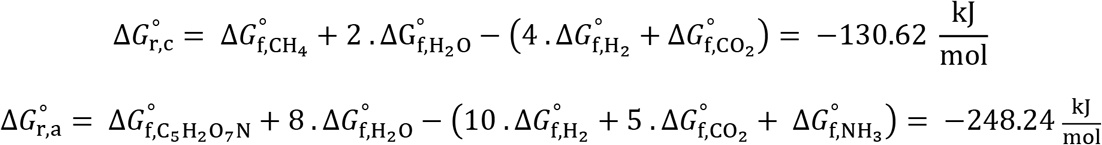

The free energy of the methanogenesis process per mole (or C-mol) of biomass formed, Δ*G*_m_, can then obtained from the following equation

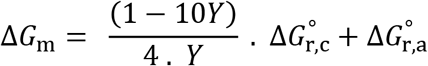

which yields for *M. smithii* and *M. ruminantium*

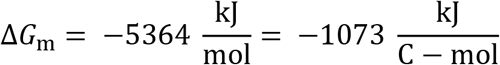

Knowing that

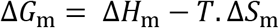

it follows that the entropic contribution to the methanogenesis process is equal to

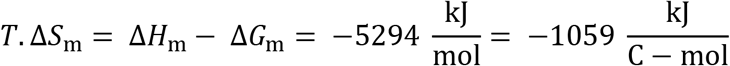

which gives, at 39°C, the following value for the entropy of the methanogenesis process per mole (or C-mol) of biomass formed

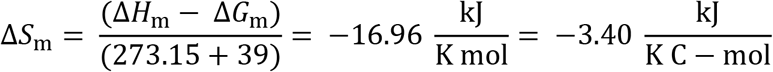

The same procedure applied to *M. formicium* yields 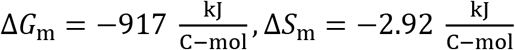

